# Impact of biosolids amendment on the soil resistome and microbiome – a greenhouse study

**DOI:** 10.1101/2024.09.30.615847

**Authors:** John Ste. Marie, Catherine Mays, Bing Guo, Tyler S. Radniecki, Joy Waite-Cusic, Tala Navab-Daneshmand

## Abstract

The spread of antibiotic resistance poses a significant challenge to public health worldwide. Wastewater treatment utilities are reservoirs of antibiotic-resistant bacteria and antibiotic resistance genes (ARGs). This study investigates the impact of biosolids amendment on the soil microbiome, resistome, virulence factors, and ESKAPE pathogens during carrot cultivation in a replicated greenhouse study. Metagenomic sequencing revealed that biosolids amendment increased the richness of microbial genera, ARGs, and virulence factors in soil. The relative abundance of ESKAPE pathogens, including *Enterococcus faecium, Staphylococcus aureus, Klebsiella pneumoniae, Acinetobacter baumanii, Pseudomonas aeruginosa*, and *Enterobacter* spp., was notably higher in biosolids-amended soils. These pathogens persisted throughout the 11-week cultivation period, raising concerns about the potential for horizontal gene transfer and the spread of antibiotic resistance. The study also identified significant co-occurrences between microbial genera and ARGs, which can suggest the possibility of the microbial taxa harboring the co-occurring ARGs. These findings highlight the importance of continued research and monitoring to ensure the safe and effective use of biosolids in agriculture.

## Introduction

The increased threat of antibiotic resistance, recently dubbed “The Silent Pandemic”, is a rapidly growing public health concern (G7 Health Ministers, 2022). Wastewater treatment utilities are one of the main recipients, reservoirs, and resources of antibiotic-resistant bacteria (ARB) and antibiotic resistance genes (ARGs) (Karkman et al., 2018; Pazda et al., 2019). Additionally, wastewater utilities receive a wide variety of antibiotics, metals, and other antimicrobials that can promote the horizontal transfer of ARGs (Anjali and Shanthakumar, 2019; Wang et al., 2020). In wastewater treatment trains, a large fraction of the ARB, ARGs, and mobile genetic elements end up in the biosolids (Munir et al., 2011).

In the United States, 31% of biosolids are used as amendments to agricultural soils to improve soil properties and provide valuable nutrients (EPA, 2023). Consequently, environments receiving biosolids have been shown to contain a large diversity of ARB (including human pathogens) and ARGs and are considered reservoirs for antibiotic resistance propagation (Berendonk et al., 2015; Cerqueira et al., 2019; D’Angelo, 2023; Kim and Cha, 2021). Additionally, biosolids contain diverse microorganisms, with large prevalences of *Proteobacteria* and *Bacteroidetes* which contain pathogenic species. Accordingly, there are concerns that biosolids land application may increase the prevalence of human pathogens in biosolids-amended soils and further propagate clinical antibiotic resistance.

Among the human pathogens of greatest concern are the ESKAPE pathogens: *Enterococcus faecium, Staphylococcus aureus, Klebsiella pneumoniae, Acinetobacter baumanii, Pseudomonas aeruginosa*, and *Enterobacter* spp. These pathogens are known for their multidrug resistance and clinical prevalence, posing a direct public health threat (Santajit and Indrawattana, 2016). ESKAPE pathogens have been detected in wastewater and receiving environments, including multidrug-resistant Acinetobacter and *Pseudomonas aeruginosa*, and methicillin-resistant *Staphylococcus aureus* (MRSA) (Kozajda and Jezak, 2020; Nishiyama et al., 2021). While recent research has reported on ESKAPE taxa in wastewater influent and effluent (Ramos et al., 2024; Raza et al., 2022), there is not a comprehensive understanding of the fate of these pathogens in environments amended with biosolids.

In a broader context, longitudinal studies have examined the impact of biosolids amendment on the soil microbiome and ARG abundance. Many of these studies rely on targeted analyses, such as qPCR, to characterize the soil resistome (Cerqueira et al., 2019; Schlatter et al., 2019; Zhang et al., 2019). While qPCR analysis provides an accurate quantitative assessment of the abundances of targeted ARGs, they are limited in their ability to examine the breadth of diversity in the resistome (Liguori et al., 2022).

To address this limitation, metagenomic approaches have been used to characterize changes in the microbiomes and resistomes in soils amended with both untreated and composted manure at the microcosm, greenhouse, and field scales (Chen et al., 2019; Guron et al., 2019; Keenum et al., 2022). These studies take advantage of the vast data acquired from metagenomic analysis and use network analysis and other emerging approaches to determine associations between the microbiome and resistome and evaluate the risk of manure amendment. Metagenomic analyses have also been successfully used to characterize other impacted soils such as those irrigated with wastewater influent, chronically exposed to heavy metals, and amended with antibiotics (Bougnom et al., 2019; Brown Liam P. et al., 2022; Salam, 2020).

While there have been some metagenomic studies on biosolids-amended soils, they are primarily focused on the biosolids themselves or the impact of biosolids treatment rather than their impact on soil microbiomes and resistomes (D’Angelo, 2023; Guo et al., 2017; Han and Yoo, 2020; Yergeau et al., 2016). Thus, more comprehensive longitudinal replicated analysis of soils amended with biosolids are needed to understand the impact of biosolids amendments on soil microbiomes and resistomes. In this study, shotgun metagenomics was used to characterize the impact of biosolids amendment on the enrichment of the microbiome, resistome, virulence factors, and ESKAPE pathogens in soils during carrot cultivation in a replicated greenhouse study. The relationships within and between the microbiome and the resistome were identified to elucidate co-occurrences between the microbiome and ARGs.

## Materials and Methods

### Sample Collection

The greenhouse study design has been described in detail previously (Mays et al., 2021). Briefly, the soil for cultivation was collected from a commercial agricultural field with no history of wastewater irrigation or biosolids amendment in the Willamette Valley area of Oregon. The soil was air-dried and passed through a 5-mm sieve. Dewatered Class B biosolids were obtained from a conventional wastewater treatment utility in Oregon that uses a conventional activated sludge process. The biosolids were transferred to the laboratory on ice and applied to triplicate pots at a ratio of 70 g per kg of soil. Similarly, triplicate pots of soil – herein referred to as pristine soil – were used as the control treatment. Four germinated carrot seeds were planted in each of the six pots and cultivated for 11 weeks, at which time they were harvested. Deionized water was used to irrigate the plants two to three times per week to maintain a soil moisture content of 80-85% total solids.

Soil core samples were collected from the bulk soil using a soil sampler probe (M.K. Rittenhouse & Sons Ltd., St. Catharines, Ontario, Canada) at the time of planting, on week 6 of the study, and at week 11 at the time of harvest. Soil samples were transferred in WhirlPak bags (Nasco, Fort Atkinson, WI) to the lab where they were homogenized by massaging and shaking for 2 min. From each sample, approximately 0.5 g of soil was stored in 50% v/v ethanol at -20 °C for microbial analysis. Genomic DNA was extracted using the FastDNA Spin Kit for Soil (MP Biomedicals, Irvine, CA). Extracted DNA was stored at -20 °C until submission for sequencing.

### DNA purification, shotgun sequencing, and quality analysis

The concentration and quality of extracted DNA were measured with a Qubit fluorometer (Thermo Fisher Scientific, Waltham, MA). To improve the quality of low-concentration samples, extracted DNA was purified with a ReliaPrep DNA Clean-up and concentration kit (Promega, Madison, WI). Purified DNA was submitted to the Center for Quantitative Life Sciences at Oregon State University. Sequencing libraries were prepared for all samples using the NexteraXT kit (Illumina, San Diego, CA), except for the initial biosolids-amended samples, which were prepared using PrepX (IntegenX, Pleasanton, CA). The three biosolids-amended samples required the PrepX kit because the BioAnalyzer results indicated DNA quality issues when using the NexteraXT kit. Paired-end (2×150 bp) sequencing reads were generated on an Illumina HiSeq 3000 (Illumina, San Diego, CA). Read quality was assessed with FastQC (Andrews, 2010). Subsequently, reads were filtered and trimmed using CutAdapt with quality thresholds of 20 and 15 for forward and reserve reads, respectively, and a minimum length of 36 (Martin, 2011). Reads were again assessed with FastQC to verify quality before subsequent analysis.

### Metagenomic Analysis

The microbial community structure was characterized using Kaiju (Menzel et al., 2016), which translates the raw metagenomic reads into amino acid sequences in all six possible reading frames and searches for the maximum exact matches in the microbial reference gene database from the National Center for Biotechnology Information (NCBI; Menzel et al., 2016). Those that are matched are then assigned to a taxon in the NCBI taxonomy.

Cleaned reads from each sample were then *de novo* assembled with MEGAHIT (D. Li et al., 2015) using a minimum k value of 59, and a maximum of 159. Assembled reads were processed with ARGs-OAP to annotate ARGs using the SARG database, a non-redundant ARG reference database integrating the Comprehensive Antibiotic Resistance Database (CARD) and the Antibiotic Resistance Database (ARDB) (Yin et al., 2018). This hierarchical classification annotated the reads at the type and subtype levels using hidden Markov models. Virulence factors (VFs) in contigs were identified through a BLAST search against the virulence factor database (VFDB), a comprehensive database containing virulence factors from various bacterial pathogens (Liu et al., 2019).

### Statistical Analysis

To assess the alpha diversity (richness and evenness) within each treatment (pristine soil and biosolids-amended soil at weeks 0, 6, and 11), the Shannon index and richness for each treatment’s microbiome (species level), resistome, and VFs were calculated in R (version 4.2.2) using the Vegan package (Oksanen, 2024). The impacts of biosolids amendment on the resistome and microbiome diversities were determined by comparing the Shannon indices of pristine samples and those amended with biosolids using the student’s t-test. To determine the differences between sample groups in the microbiome (at the genus level), resistome, and VFs, the beta diversity was determined using the Bray-Curtis dissimilarity and the Vegan package. The variances between treatment groups were determined using the permutational multivariate analysis of variance (PERMANOVA) test Adonis in the Vegan package, which reduces the dimensionality of the data to identify statistically significant differences between biosolids-amended and pristine soil. Hierarchical cluster analysis was used to create heatmaps of the microbial phyla and genera (present above 0.1%) as well as ARGs across samples using the Vegan package. In the heatmaps, dendrograms were generated using Euclidean distances.

The relative abundance of ARGs at the subtype level and microbial taxa at the genus level were aggregated. Data was filtered to include microbial genera and ARGs present in at least three samples. To determine co-occurrences within and between the microbiome and resistome, pair-wise Spearman correlations were determined. Statistically correlated (*p* < 0.01) ARG subtypes and microbial genera were visualized as a network with the R package igraph (Csardi and Nepusz, 2006).

## Results and Discussion

### Metagenomic Sequencing Data

An average of 18 ± 4.0 (± indicates the standard error) million and 20 ± 2.8 million reads were generated from the pristine and biosolids-amended soils, respectively (Table S1). Trimmed reads had an average of 17.6 ± 4.0 million clean reads for pristine soils and 19.8 ± 2.9 million clean reads for biosolids-amended soils. Read assembly resulted in a total of 4,641,285 contigs with an average of 143,670 ± 21,757 contigs in pristine soils compared to an average of 403,955 ± 25,612 contigs in biosolids-amended samples (Table S2). The average N50 for all samples was 583 ± 28.8, with significantly higher values in biosolids-amended samples (average 657 ± 31.5) compared to pristine samples (average 457 ± 19.7; one-sided t-test *p* < 0.01; Table S2). During weeks 6 and 11, one pristine sample had too little DNA to adequately sequence and was removed from further analysis. The average number of mapped reads to microbial taxa for each time point and condition can be found in supplementary Table S1.

### Microbial Community Diversity and Profile

The impact of biosolids amendment on the microbiome diversity was identified using alpha (Figure 1) and beta (Figure 2) diversity indices. Including all time points, there were no significant differences between the Shannon indices of microbial genera in the pristine (4.5 ± 0.2) and biosolids-amended (4.4 ± 0.1) soils (one-sided t-test, *p* > 0.05; Figure 1a). These Shannon indices are generally lower than reports on agricultural soils from Arizona and California which ranged between 6.0-7.3 but were comparable to those from biosolids from conventional wastewater treatment facilities which ranged from 3.4 to 5.3 (Ma et al., 2016; Wolters et al., 2022).

**Figure 1.**
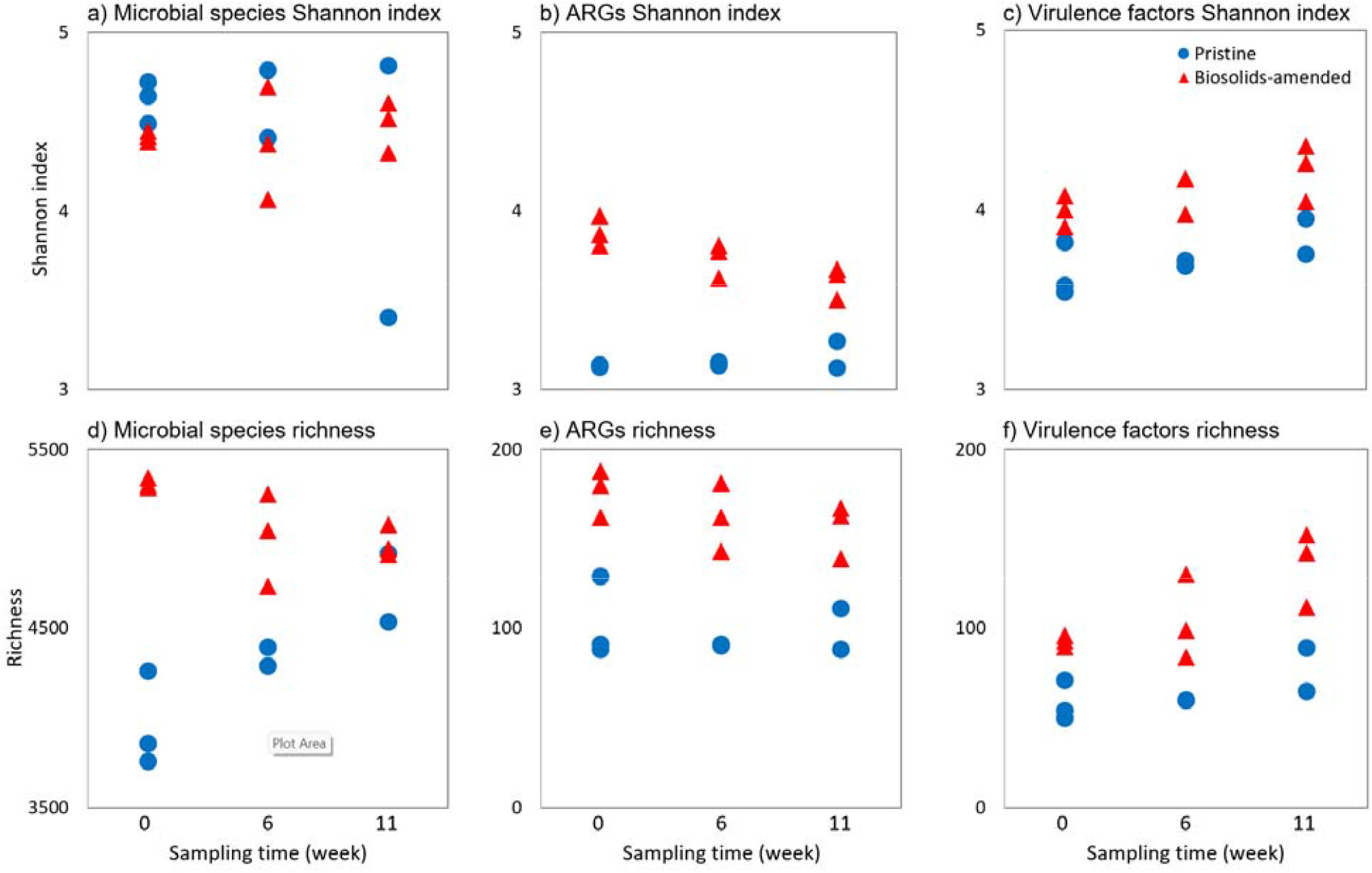
Alpha diversity indices (a-c) Shannon index and (d-f) richness for (a) and (d) microbial species, (b) and (e) antibiotic resistance genes (ARGs), and (c) and (f) virulence factors in pristine and biosolids-amended soils at weeks 0, 6, and 11 in a replicated greenhouse study.

**Figure 2.**
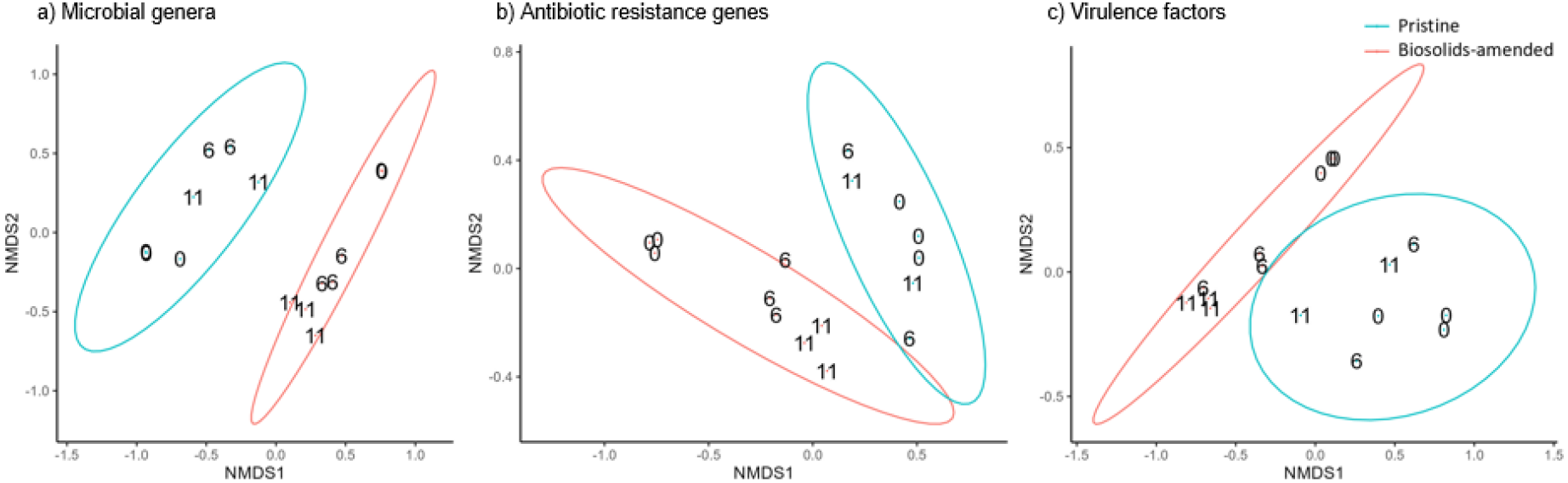
Ordination of the Bray-Curtis dissimilarity between the (a) microbial genera, (b) antibiotic resistance genes, and (virulence factors) of pristine and biosolids-amended soils. Labels indicate the sampling time (*i*.*e*., weeks 0, 6, and 11) and the ellipses denote the 95% confidence level for each treatment condition (*i*.*e*., pristine and biosolids-amended soils).

The richness of microbial species, however, was statistically larger in biosolids-amended soils (5,310.7 ± 16.5) compared to pristine (3,959.7 ± 153.3) samples (one-sided t-test, *p* < 0.001; including all the time points; Figure 1d). However, the difference in richness declined from the time of the amendment (5,310 ± 16 in biosolids-amended soils vs. 3,960 ± 153 in pristine) to the time of harvest (4,977 ± 51 in biosolids-amended vs. 4,727 ± 191 in pristine soils). This decrease in the genera richness may be attributable to the bacteria unique to the biosolids not naturalizing into the soil and dying out over time. Similar results of the gradual return from microbiome enrichment in biosolids-amended soils to comparable conditions with control soil have been previously documented (Zhang et al., 2018).

The relative abundances of microbial phyla and genera in each sample were determined to understand the impact of biosolids amendment and cultivation time on the microbial community structure. Across all sample types, there was an average of 35.3 ± 1.7% of reads unmapped to any phyla in the database, which represented the largest relative abundance of reads for all samples. Of the mapped reads, the most abundant phylum in both pristine and biosolids-amended samples was *Proteobacteria* throughout the cultivation period (Figure S1). *Bacteroidetes* was the second most abundant phyla in all samples at week 0. Previous studies have also observed these phyla to be dominant in both pristine soils and gut microbiomes (Fierer et al., 2007; Lozupone et al., 2012).

Biosolids-amended samples (*n* = 9) contained a statistically larger enrichment of the phylum *Bacteroidetes* (14.5 ± 1.1%) as compared to pristine soil samples (*n* = 7; 6.3 ± 1.0%; one-sided t-test, *p* < 0.01). *Bacteroidetes* remained the second most abundant phylum in biosolids-amended samples for the duration of the study, while in pristine samples *Actinobacteria* was the second most dominant phylum during weeks 6 and 11. Because *Bacteroidetes* are relatively prevalent in both human guts and natural soils, the enrichment of this phylum after soil amendment throughout the cultivation period could be due to their ability to secrete a wide range of carbohydrate-active enzymes capable of targeting diverse and complex glycans present in the soil (Larsbrink and McKee, 2020).

Analyses at the genera level identified *Massilia, Ralstonia, Mucilaginibacter, Nocardioides, Streptomyces, Sphingomonas*, and *Janthinobacterium* (clusters A and C) as the dominant genera in pristine soils at Week 0 (Figure 3). Biosolids amendment, however, altered the microbial communities so that the dominant genera at week 0 included *Dechloromonas, Sulfuritalea, Flavobacterium, Candidatu, Accumulibacter*, and *Pseudomonas* (Clusters E and F). The enrichment of *Dechloromonas* in the biosolid amended soil is consistent with the prevalence of these genera in biosolids and its association with the human microbiome (Newton et al., 2015; Su et al., 2017).

**Figure 3.**
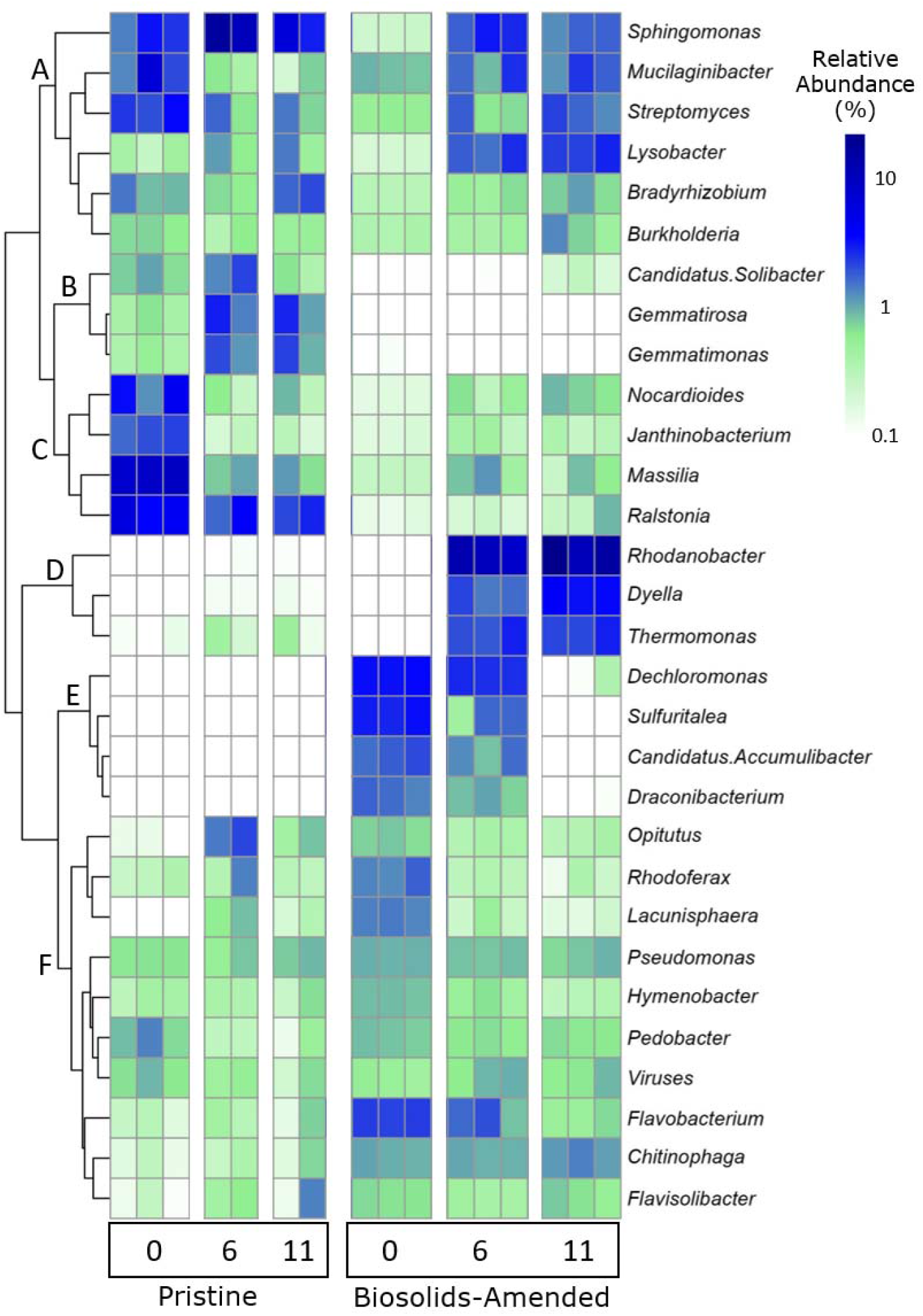
Relative abundance (%) of the top 30 most abundant genera detected in at least three samples in pristine and biosolids-amended soil samples collected at weeks 0, 6, and 11 in a greenhouse study. The dendrogram was generated using Euclidean distance.

By the end of the study (*i*.*e*., Week 11), the abundance of these notable genera became more similar between pristine and biosolids-amended soils. The main differentiating genera between the two treatments at Week 11 were *Ralstonia, Gemmatirosa*, and *Gemmatimonas* (larger abundances in pristine soils) and *Rhodanobacter, Dyella*, and *Thermomonas* (larger abundances in biosolids-amended soils). These distinguishing genera between the two treatments at different cultivation points can explain the statistical dissimilarities between the pristine and biosolids-amended soils (PERMANOVA, *p* < 0.001; Figure 2a).

Thus, these findings demonstrate that the amendment of biosolids to soil increased the richness of microbial species and resulted in statistically larger dissimilarities in microbial genera between the pristine and biosolids-amended soil. Additionally, these findings demonstrate the statistically large impact of biosolids amendment on the soil microbial community (clusters A, C, E, and F). However, with cultivation time, microbial communities become more similar in terms of clusters A, E, and F and more dissimilar in clusters B and D. Therefore, the amendment of biosolids resulted in a lasting impact of over 11 weeks on the soil microbial community structure.

### Antibiotic Resistance Genes Profile and Diversity

The impacts of biosolids amendment on the soil resistome were quantified using alpha and beta diversity indices. Biosolids-amended soils had a significantly higher diversity (Figure 1b) and richness (Figure 1e) of ARG subtypes than pristine soil samples (one-sided t-test, *p* < 0.001). For example, biosolids-amended samples contained 165 unique subtypes (genotypes) of ARGs while the pristine soils only contained 98 unique subtypes. The enrichment of ARGs in soils amended with biosolids has been previously documented (Bondarczuk et al., 2016; Markowicz et al., 2021; Yang et al., 2018; Zhang et al., 2018).

Additionally, the resistome of the pristine and biosolids-amended soils was statistically distinct (PERMANOVA, *p* < 0.001, Figure 2b). Higher diversity of ARGs can confer ecological resilience to microbial communities, allowing them to adapt to changes in environmental conditions, including antibiotic exposure (Klümper et al., 2024). However, higher diversity also raises concerns about the potential for the horizontal gene transfer of these resistance genes, which could contribute to the spread of antibiotic resistance in agricultural settings (Manyi-Loh et al., 2018).

Resistance to 19 different classes of antibiotics as well as multidrug resistance classes was observed (Figure S3). The most prevalent type of resistance in all but one sample was multidrug resistance, with an average prevalence across all samples of 0.063 ± 0.006 gene copies/16S rRNA. The pristine soil sample S014, on Week 0, harbored vancomycin resistance genes (0.051 gene copies/16S rRNA) as the most abundant antibiotic class followed by multidrug resistance (0.046 gene copies/16S rRNA). Across all samples, the next most abundant classes of resistance were to vancomycin (0.021 ± 0.004 gene copies/16S rRNA), bacitracin (0.018 ± 0.002 gene copies/16S rRNA), fosmidomycin (0.013 ± 0.002 gene copies/16S rRNA), and sulfonamides (0.009 ± 0.002 gene copies/16S rRNA).

Biosolids amendment increased the abundance of resistance genes belonging to several classes of antibiotics including sulfonamide, tetracycline, fosmidomycin, and macrolides, an impact that persisted throughout the 11 weeks. Other studies have reported similar enrichments of tetracycline, sulfonamide, and beta-lactam resistance genes after biosolids soil amendment (Hung et al., 2022; Lau et al., 2017; Qin et al., 2022). The only type of resistance that was more abundant in pristine soils compared to the biosolids-amended soils was resistance to vancomycin with an average relative abundance of 0.035 ± 0.006 gene copies/16S rRNA copy in pristine samples and only 0.010 ± 0.003 gene copies/16S rRNA copy in biosolids. These results demonstrate that in addition to the impact on resistome diversity, biosolids amendment changes the classes of resistance genes in soils.

The temporal impact of biosolids amendments to the soil resistomes was quantified through the relative abundances of ARG subtypes. At the beginning of the study (*i*.*e*., Week 0), the most abundant ARG subtypes in pristine soils included the vancomycin resistance gene *van*R, followed by *bac*A and the multidrug resistance gene *mdt*B (Figure 4). In biosolids-amended soils, at the time of amendment (*i*.*e*., Week 0), the most abundant subtype was *bac*A, conferring resistance to bacitracin, followed by the sulfonamide resistance gene *sul*1 and the multidrug resistance gene *qac*E_Δ_1. At Week 11, there was little change in the resistome of the pristine soils with *van*R and *bac*A remaining the most dominant ARG subtypes. In biosolids-amended soils, however, the most abundant ARG subtypes at Week 11 were the multidrug resistance gene *mex*F, the fosmidomycin resistance gene *ros*A, and a multidrug transporter gene.

**Figure 4.**
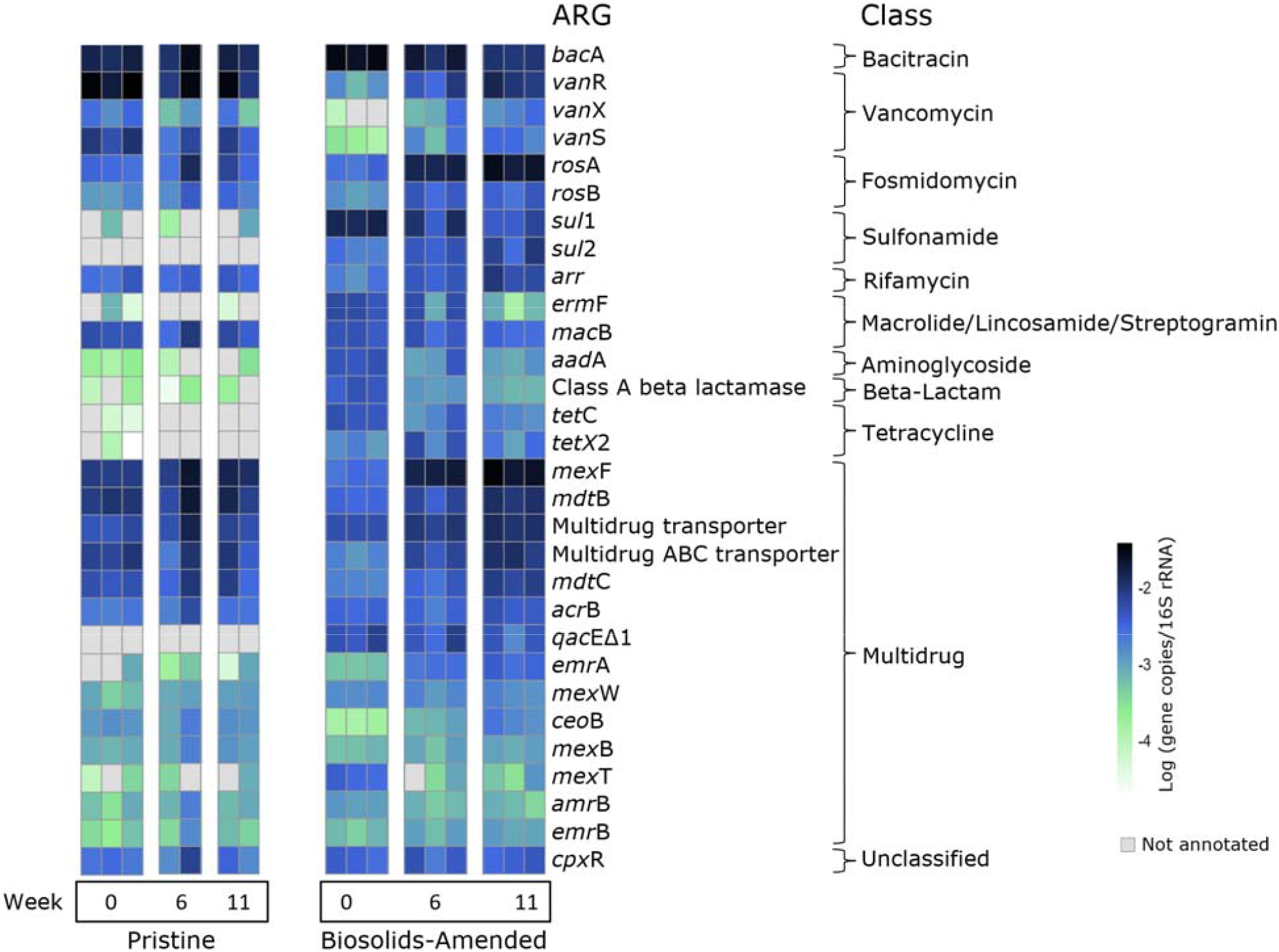
Heatmap of the 30 most abundant antibiotic resistance gene (ARG) subtypes detected in at least three samples in pristine and biosolids-amended soils collected at weeks 0, 6, and 11 in a greenhouse study. Abundance is shown in log (gene copies/16S rRNA). ARG classes are shown to the right.

The difference in ARG subtype relative abundances between the two treatments at different cultivation points can explain the statistical dissimilarities between the pristine and biosolids-amended soils (PERMANOVA, *p* < 0.001; Figure 2b). Additionally, even though the ARG subtype relative abundances change over time for the biosolids-amended soils, they were still different than the ARG subtypes found in the pristine soils. Thus, these findings demonstrate the statistically large impact of biosolids amendment on the soil resistome, with some of these impacts persisting throughout the 11-week cultivation period. Similar results of long-lasting persistence of biosolids-derived ARG subtypes following land application has been previous documented (Berendonk et al., 2015; Cerqueira et al., 2019; Kim and Cha, 2021).

### Virulence Factors Profile and Diversity

A total of 285 VFs were annotated across all samples. Including all time points, biosolids amendment resulted in significantly higher alpha diversity of VFs measured by Shannon index (4.11 ± 0.05) compared to pristine soils (3.72 ± 0.05; one-sided t-test, *p* < 0.001, Figure 1c). The soil richness of VFs was also significantly higher in biosolids-amended (110.9 ± 8.2) compared to pristine samples (64.1 ± 4.9; one-sided t-test, *p* < 0.001, Figure 1f). The difference in VFs between pristine and biosolids-amended soils (*i*.*e*., treatments) was statistically larger than the differences observed within treatments (PERMANOVA, *p* < 0.001; Figure 2c).

The higher alpha and beta diversities in the biosolids-amended soils suggest that the pathogens possess a wider array of mechanisms and have a higher potential to cause diseases. A diverse range of virulence factors can lead to complex interactions within microbial communities. This complexity can influence disease outcomes, the severity of infections, and the effectiveness of treatments or interventions (Martínez José L. and Baquero Fernando, 2002).

### ESKAPE Pathogens Profile

Across all time points (*i*.*e*., Weeks 0, 6, and 11) biosolids-amended soils (0.015 ± 0.003%) had higher relative abundances of all six ESKAPE pathogens compared to pristine soils (0.007 ± 0.001%; *p <* 0.01) (Figure S2). At the time of planting, biosolids-amended soils showed higher abundances of each ESKAPE pathogen except for *Staphylococcus aureus*, which had comparable relative abundance in pristine (0.0033%) and biosolids-amended (0.0035%) soils. At the time of harvest, biosolids amendment still had significantly higher enrichment of the six ESKAPE pathogens.

Relative to pristine samples at harvest, biosolid-amended soils had 232% more *Enterococcus faecium* (one-sided t-test, *p* = 0.009), 55.8% more *Staphylococcus aureus* (*p* = 0.088), 57.5% more *Klebsiella pneumoniae* (*p* = 0.005), 241% more *Acinetobacter baumanii* (*p* < 0.001), 164% more *Pseudomonas aeruginosa* (*p* < 0.001), and 38.6% more *Enterobacter* spp. (*p* = 0.002). Therefore, biosolids amendment to soil increased the abundance of six pathogens with great threat to human health with lasting impact throughout the 11-week cultivation period of our study. However, while ESKAPE pathogens were present and enriched in our study, they remained at a low relative abundance in soil (0.007 ± 0.001% in pristine and 0.015 ± 0.003% in biosolids-amended soils) similar to previously reported data (Schlatter et al., 2019).

The increase in the alpha diversity indices (Shannon index and richness) of ARG subtypes (Figures 1b and e; *p* < 0.001) and virulence factors (Figures 1c and f; *p* < 0.001) as well as the relative abundance of ESKAPE pathogens (Figure S2; *p* < 0.01) in the biosolids-amended soil is of concern. This demonstrates the potential for high-risk pathogenic bacteria to acquire virulence factors and ARGs following the amendment of biosolids in agricultural soils via opportunistic horizontal gene transfer. For example, *Pseudomonas aeruginosa*, an ESKAPE pathogen, has been shown to be responsible for 11% of virulence gene expression in biosolids-amended soils (D’Angelo, 2023). ESKAPE pathogens can acquire ARGs through horizontal gene transfer as they often host ARG-laden mobile elements such as plasmids and prophages (Das et al., 2022). Similarly, other pathogens can horizontally transfer clinically relevant ARGs to environmental bacteria when exposed to selective pressures, making biosolids-amended soils a potential reservoir for high-risk ARGs and pathogens (Bondarczuk et al., 2016).

### Associations between the Microbiome and Resistome

Co-occurrence analyses within and between the microbial genera and ARGs (Figure 5) identified statistically large (*p* < 0.01 and Spearman coefficient > 0.8) associations (*i*.*e*., edges) within and between the microbial genera and ARGs (*i*.*e*., nodes). In pristine soil, eight clusters of microbial genera with at least three nodes were identified, while all other observed associations contained only two nodes (Figure 5a). One large cluster included six microbial genera (*Solitalea, Sulfuriferula, Algoriphagus, Rufibacter, Saprospira*, and *Echinicola*) and the tetracycline resistance gene *tet*O. Of these six microbial genera, five belonged to the Bacteroidota phylum while the other one (*Sulfuriferula*) belonged to Pseudomonadota. A previous study on target ARGs and amplicon-sequenced microbial communities reported co-occurrences of tetracycline resistance genes (*e*.*g*., *tet*G and *tet*X) with microbial genera belonging to the Bacteroidota phylum (*e*.*g*., *Parapedobacter, Sphingobacteriaceae, Sphingobacterium*, and *Flavobacterium*) following the land application of sludge composts (Zhang et al., 2018).

**Figure 5.**
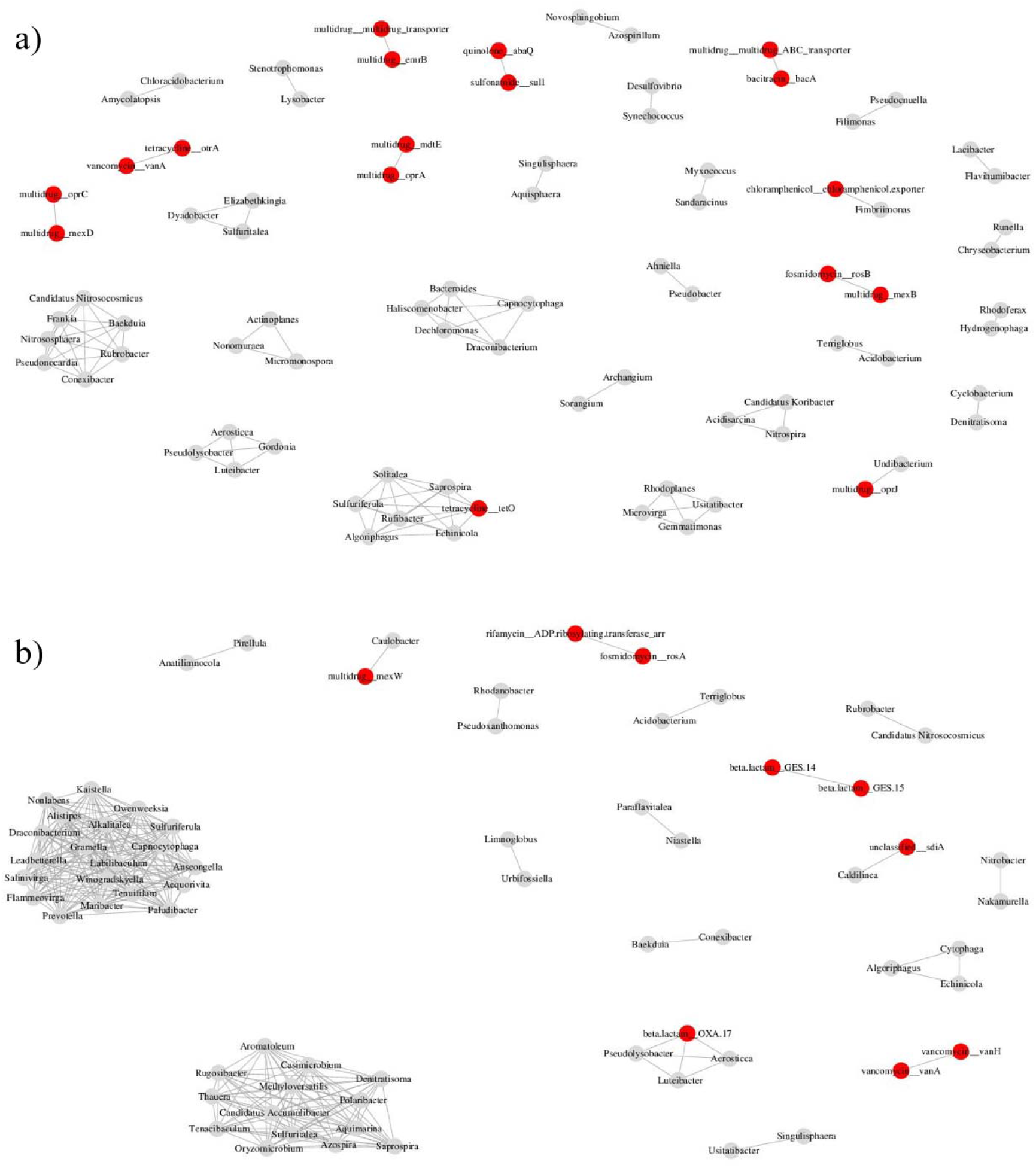
Associations between the microbiome (at the genus level present in at least three samples) and resistome in (a) pristine and (b) biosolids-amended soils in collected at weeks 0, 6, and 11 in a greenhouse study. All associations (shown by connections) represent significant Spearman correlations at *p* < 0.01. Grey symbols represent the microbial genera and red symbols indicate antibiotic resistance genotypes.

Additionally, in pristine soils, the co-occurrence associations of two Proteobacteria genera (*Undibacterium* and *Fimbriimonas*) with the multidrug resistance gene *oprJ* and a chloramphenicol transport gene, respectively, were observed (Figure 5a). Three multidrug resistance pairs were identified including *mex*D and *opr*C, *erm*B and a multidrug transport gene, and *opr*A and *mdt*E. Multidrug resistance prevalence was also associated with resistance to bacitracin (*bac*A) and fosmidomycin (*ros*B). Tetracycline resistance gene, *otr*A was associated with the prevalence of *van*A gene resisting vancomycin. The sulfonamide resistance *sul*1 prevalence clustered with the quinolone resistance *aba*Q. Despite these associations, none of the identified taxa in the clusters were abundant at levels greater than 0.1% in pristine samples.

In biosolids-amended samples, two large and interconnected clusters of microbial genera were found (Figure 5b). In one of these clusters, all genera were from the Bacteroidota phylum except *Sulfuriferula* from the Pseudomonadota phylum. The second large microbial cluster contained genera from the Bacteroidota and Pseudomonadota phyla as well as *Casimicrobium* from the Firmicutes phylum. The Bacteroidota phylum is the primary degrader of complex carbohydrates found in soils (Larsbrink and McKee, 2020), therefore their presence is likely originated from the soil microbiome. *Sulfuriferula* (from Pseudomonadota phylum) are sulfur-oxidizing chemolithoautotrophs – organisms that derive energy from chemical reactions of reduced compounds of mineral origin, making soil the preferred environment for them (Kojima et al., 2020).

In addition to the two large clusters of microbial genera, two other clusters of genera contained more than three nodes, one of which included a beta-lactam resistance gene *oxa*17. Two pairs of genera and ARG subtypes co-occurrences were identified including the multidrug efflux pump *sdi*A and *Caldilinea* as well as the multidrug resistance gene *mex*W and *Caulobacter*. Two beta-lactam resistance genes *ges*14 and *ges*15 and two vancomycin resistance *van*A and *van*H also co-occurred in biosolids-amended soils. Additionally, the fosmidomycin resistance *ros*A co-occurred with the rifamycin resistance *adp*. While many of the genera identified to have co-occurrences had relatively low abundance (< 0.1%) in biosolids-amended soils, the genera *Caldilinea* did represent 0.75% of the microbial community (Figure 5b).

This increase in microbiome network connectivity following biosolids land application has been previously observed (Price et al., 2021). This current study identified multiple co-occurrences of ARGs (with resistances to tetracyclines, beta-lactams, chloramphenicol, and multidrugs) and microbial genera in pristine and biosolids-amended soils. These co-occurrences have been attributed to the possibility of the microbial taxa harboring the co-occurring ARGs due to similar abundance trends in different environmental reservoirs (Forsberg et al., 2014; B. Li et al., 2015). The observed co-occurrence associations identified by network analysis need to be further validated using targeted approaches.

## Conclusions

This study comprehensively determines the soil microbiome, resistome, virulence factors, and the ESKAPE pathogens impacted by biosolids amendment in a controlled and replicated greenhouse setting. The results from this study demonstrate the persistent impact that biosolids soil amendment has on the receiving soil environment. Biosolids amendment increased the richness of microbial genera, antibiotic resistance genes (ARGs), and virulence factors. In addition, the abundance of the ESKAPE pathogens increased after the amendment of biosolids. While increased microbiome diversity in agricultural soils can suggest ecosystem resiliency and stability resulting in improved soil fertility and crop yield, when paired with increased richness of the resistome, it can lead to increased potential for the horizontal gene transfer of ARGs. The enrichment of the resistome, alongside ESKAPE pathogens and virulence factors following biosolids amendment, indicates the risk of introducing and naturalizing clinically relevant antibiotic-resistant pathogens into the native soil environment. This could result in the proliferation of these pathogens in any future agricultural applications of the soil, such as crop cultivation.

## Acknowledgment

This work was supported by the USDA National Institute of Food and Agriculture, Agricultural and Food Research Initiative Competitive Program, Agriculture Economics and Rural Communities, grant number: 2018-67017-27631.

## Data availability

All sequenced reads can be found in the National Center for Biotechnology Information (NCBI) Sequence Read Archive under Bioproject accession number PRJNA1049878.

