## Supplementary Information for "Impact of biosolids amendment on the soil resistome and microbiome – a greenhouse study"

**Table S1**. The number of raw and trimmed reads and the number of reads annotated by Kaiju and ARGs-OAP in all samples (Menzel et al., 2016, Yin et al., 2018a). Treatments include pristine soil (without any amendment) and biosolids-amended soil. Soil core samples were collected during weeks 0, 6, and 11 during carrots cultivation in a replicated greenhouse study.

| Treatment | Week | Sample ID | Raw | Trimmed | Kaiju  annotated reads | ARGs-OAP annotated reads |
| --- | --- | --- | --- | --- | --- | --- |
| Pristine | 0 | S014 | 13,342,500 | 12,998,034 | 69,376 | 56,202 |
|  |  | S015 | 15,698,695 | 15,389,999 | 81,894 | 63,256 |
|  |  | S016 | 21,484,135 | 21,081,486 | 124,885 | 98,109 |
|  | 6 | S007 | 16,031,417 | 15,957,118 | 144,101 | 77,829 |
|  |  | S012 | 17,982,061 | 17,019,522 | 139,248 | 74,609 |
|  | 11 | S008 | 24,982,893 | 24,874,784 | 243,950 | 118,438 |
|  |  | S013 | 16,534,041 | 16,165,375 | 202,237 | 59,865 |
| Biosolids-amended | 0 | S009 | 20,891,440 | 20,399,186 | 470,370 | 68,434 |
|  |  | S010 | 21,000,859 | 20,630,512 | 484,753 | 69,177 |
|  |  | S011 | 16,594,751 | 15,964,463 | 449,036 | 56,889 |
|  | 6 | S001 | 24,739,104 | 24,537,511 | 492,343 | 114,036 |
|  |  | S003 | 16,019,127 | 15,945,296 | 253,815 | 62,976 |
|  |  | S005 | 20,388,738 | 20,303,184 | 358,489 | 94,571 |
|  | 11 | S002 | 19,613,886 | 19,418,974 | 372,837 | 104,406 |
|  |  | S004 | 22,935,428 | 22,868,089 | 428,553 | 117,903 |
|  |  | S006 | 18,080,416 | 18,020,637 | 325,398 | 91,760 |

**Table S2**. Assembly statistics for all samples including the number of contigs, total base pairs, minimum, maximum, and average length, and average N50. Treatments include pristine soil (without any amendment) and biosolids-amended soil. Soil core samples were collected during weeks 0, 6, and 11 during carrots cultivation in a replicated greenhouse study.

| Treatment | Week | Sample ID | Sample label | Contigs | Total base pair | Min length | Max length | Avg length | N50 |
| --- | --- | --- | --- | --- | --- | --- | --- | --- | --- |
| Pristine | 0 | S014 | 1-B-0 | 69,376 | 36,069,801 | 200 | 69,345 | 519 | 495 |
|  |  | S015 | 2-B-0 | 81,894 | 43,445,440 | 200 | 69,345 | 530 | 514 |
|  |  | S016 | 1-C-0 | 124,885 | 61,625,460 | 200 | 69,345 | 493 | 454 |
|  | 6 | S007 | 1-B-6 | 144,101 | 82,340,250 | 200 | 106,282 | 571 | 565 |
|  |  | S012 | 1-A-6 | 139,248 | 58,766,287 | 204 | 19,033 | 422 | 392 |
|  | 11 | S008 | 1-E-11 | 243,950 | 118,454,278 | 200 | 61,202 | 485 | 465 |
|  |  | S013 | 1-F-11 | 202,237 | 110,243,778 | 200 | 92,298 | 545 | 526 |
| Biosolids-amended | 0 | S009 | 3-C-0 | 470,370 | 356,388,703 | 200 | 445,550 | 757 | 791 |
|  |  | S010 | 3-B-0 | 484,753 | 372,221,476 | 200 | 557,991 | 767 | 803 |
|  |  | S011 | 4-C-0 | 449,036 | 300,239,910 | 200 | 240,962 | 668 | 660 |
|  | 6 | S001 | 3-B-6 | 492,343 | 338,804,060 | 200 | 695,968 | 688 | 710 |
|  |  | S003 | 3-C-6 | 253,815 | 160,394,571 | 200 | 245,261 | 631 | 626 |
|  |  | S005 | 3-A-6 | 358,489 | 196,911,151 | 200 | 58,657 | 549 | 523 |
|  | 11 | S002 | 3-E-11 | 372,837 | 223,773,173 | 200 | 85,826 | 600 | 623 |
|  |  | S004 | 3-D-11 | 428,553 | 235,763,690 | 200 | 49,082 | 550 | 522 |
|  |  | S006 | 3-F-11 | 325,398 | 210,181,296 | 200 | 59,778 | 645 | 659 |

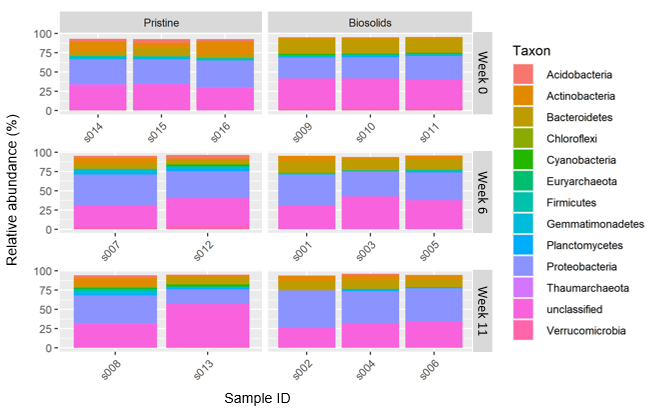

**Figure S1.** Relative abundance (%) of the dominant phyla in each sample. Samples are grouped horizontally by treatment and vertically by week of sampling. Treatments include pristine soil (without any amendment) and biosolids-amended soil. Soil core samples were collected in weeks 0, 6, and 11 during carrots cultivation in a replicated greenhouse study.

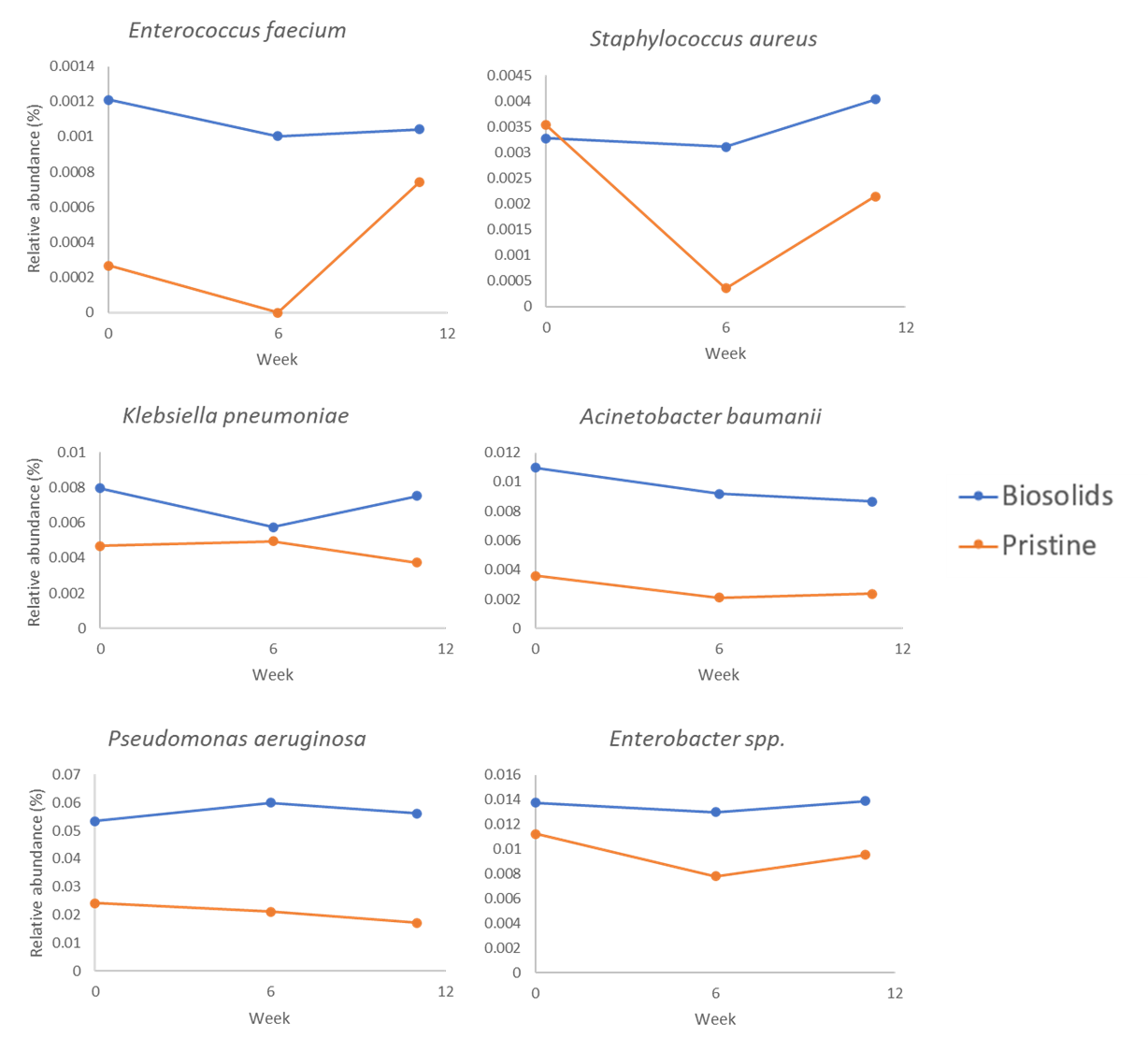

**Figure S2** Relative abundance of ESKAPE Pathogens (Enterococcus faecium, Staphylococcus aureus, Klebsiella pneumoniae, Acinetobacter baumanii, Pseudomonas aeruginosa, and Enterobacter spp.) in biosolids-amended and pristine soils.

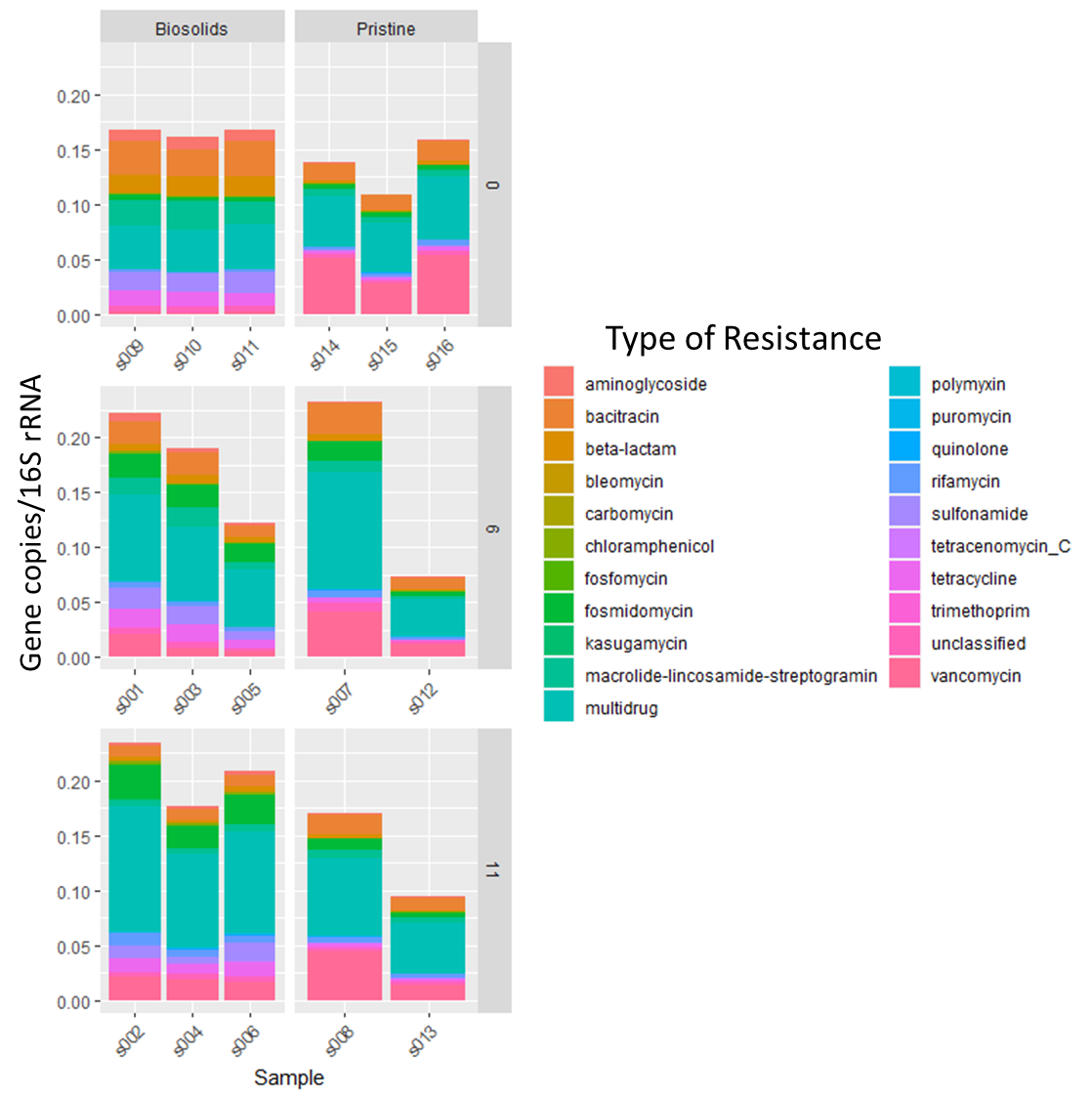

**Figure S3** Most abundant antibiotic resistance gene types (class of antibiotics resisted).
